# CD95 expression in triple negative breast cancer blocks induction of an inflammatory state through differential regulation of NF-κB Signaling

**DOI:** 10.1101/2021.04.07.438830

**Authors:** Jean-Philippe Guégan, Justine Pollet, Christophe Ginestier, Emmanuelle Charafe-Jauffret, Marcus E. Peter, Patrick Legembre

## Abstract

CD95L is expressed by tumor-infiltrating lymphocytes to eliminate CD95-expressing tumor cells and thereby CD95 loss by tumor cells is often considered as a consequence of an immunoediting process. Nonetheless CD95 expression is maintained in most triple negative breast cancers (TNBCs), and we recently reported that CD95 loss in TNBC cells triggers the induction of a pro-inflammatory program promoting the recruitment of cytotoxic NK and CD8+ T-cells and impairing tumor growth. Using a comprehensive proteomic approach, we have identified two yet unknown CD95 interaction partners, Kip1 ubiquitination-promoting complex protein 2 (KPC2) and p65. KPC2 contributes to the partial degradation of p105 (NFκB1) and the subsequent generation of p50 homodimers, which transcriptionally represses pro-inflammatory NF-κB-driven gene expression. Mechanistically, KPC2 directly interacts with the C-terminal region of CD95 and links the receptor to RelA (p65) and KPC1, the catalytic subunit of the KPC complex that acts as E3 ubiquitin-protein ligase promoting the partial degradation of p105 into p50. Loss of CD95 in TNBC cells releases KPC2, limiting the formation of the NF-κB inhibitory homodimer complex (p50/p50), promoting NF-κB activation and the production of pro-inflammatory cytokines including CSF1, CSF2, CXCL1 and IL1 members, known to promote recruitment and differentiation of certain adaptive and innate immune effector cells.

## Introduction

Among women, breast cancer (BC) is the most common cause of cancer, and the second leading cause of cancer death (DeSantis *et al*, 2016). BC is a heterogeneous disease whose molecular classification has been significantly improved to distinguish luminal A and B expressing hormonal receptors including estrogen (ER) and/or progesterone receptors (PR), basal/triple negative breast cancer (TNBC), and human epidermal growth factor receptor 2 (HER2)-like tumors. This molecular taxonomy is clinically relevant with basal/TNBC patients presenting the poorest clinical outcome with no targeted therapies available when compared to other molecular subtypes. TNBCs progress rapidly and generate metastases that remain the major cause of cancer-related mortality in these patients (Christofori, 2006). TNBC presents a major intratumoral heterogeneity that contributes to therapy failure and disease progression. The origin of this cellular heterogeneity is mainly explained by a hierarchical organization of tumor tissues where several subpopulations of self-renewing breast cancer stem cells (bCSCs) sustain the long-term oligoclonal maintenance of the neoplasm (Kreso & Dick, 2014).

CD95 (Fas/APO-1/TNFRSF6) belongs to the tumor necrosis factor (TNF) receptor family and has long been considered only as a death receptor (Peter *et al*, 2015). However, more recent data highlighted that this receptor can also induce non-apoptotic signaling pathways involved in physiological processes (Desbarats *et al*, 2003; Desbarats & Newell, 2000) or in the progression of auto-immune (O’ Reilly *et al*, 2009; Poissonnier *et al*, 2016; Tauzin *et al*, 2011) and cancer disorders (Barnhart *et al*, 2004; Kleber *et al*, 2008). CD95L (CD178), the ligand of CD95, is a member of TNF receptor superfamily and is expressed by activated T lymphocytes and NK cells to kill infected and transformed cells through cell-to-cell contact (Suda *et al*, 1993). CD95L extracellular domain can also be cleaved by metalloproteases (Fouque *et al*, 2014) to release a soluble CD95L (s-CD95L) into the bloodstream. Unlike membrane-bound CD95L (m-CD95L), s-CD95L fails to trigger cell death but instead contributes to aggravating inflammation in chronic inflammatory disorders such as systemic lupus erythematosus (SLE) (O’ Reilly *et al*., 2009; Tauzin *et al*., 2011) and cancers (Barnhart *et al*., 2004; Hoogwater *et al*, 2010; Kleber *et al*., 2008; Malleter *et al*, 2013). The intracellular death domain (DD) of CD95 serves as a docking platform to trigger cell death. Upon binding of m-CD95L, CD95 recruits the adaptor protein Fas Associated Death Domain (FADD) through homotypic interactions via their respective DD. FADD in turn aggregates the initiator caspase-8 and −10. The CD95/FADD/caspase complex called death-inducing signaling complex (DISC) contributes to the induction of the apoptotic program (Kischkel *et al*, 1995). S-CD95L fails to induce formation of the DISC but rather induces the formation of a non-apoptotic complex termed motility-inducing signaling complex (MISC) implementing a Ca^2+^ response, which promotes migration of cancer cells (Kleber *et al*., 2008; Malleter *et al*., 2013; Tauzin *et al*., 2011). The CD95-mediated Ca^2+^ response relies on a juxtamembrane region (amino acid residues 175 to 210) that we designated as calcium-inducing domain (CID). While the DD (amino acid residues 210 to 303) is involved in cell death, the biological roles of the last 15 aa of CD95 (amino acid residues 303 to 319) remain largely unknown. The protein tyrosine phosphatase FAP-1 (Sato *et al*, 1995) or Dlg1 (Gagnoux-Palacios *et al*, 2018) can interact with this carboxy-terminal region and inhibit cell death, through molecular mechanisms that remain to be elucidated. Interestingly, although CD95L-expressing immune cells might edit tumor cells by sparing cancer cells expressing low CD95 level at their plasma membrane (Strasser *et al*, 2009), complete loss of CD95 is deleterious to tumor growth (Chen *et al*, 2010). TNBCs keep the highest amount of CD95 as compared to other breast cancers (Blok *et al*, 2017), and we recently showed that loss of CD95 in TNBC reprograms the immune landscape, engendering a pro-inflammatory response and in turn promoting a natural killer (NK) and CD8^+^ T-cell anti-tumor response. The molecular mechanism responsible for the recruitment and activation of these cells remains to be elucidated.

Herein, we established that CD95 directly binds KPC2 (Kip1 ubiquitylation-promoting complex 2), which together with KPC1 forms the ubiquitin ligase KPC, and causes the partial degradation of p105 into p50. Loss of CD95 in TNBC results in the release of KPC1/KPC2 from the receptor, reducing the amount of p50 and favoring the formation of p50/p65 heterodimers. In these otherwise CD95 signaling deficient cells, this promotes the induction of a pro-inflammatory NF-κB signaling pathway resulting in the release of a set of chemo- and cytokines known to recruit multiple immune effector cells including NK and CD8+ T cells.

## Results and Discussion

### Loss of CD95 in TNBC cells activates an inflammatory signaling pathway and prevents migration of CSCs

To address the question of whether the loss of CD95 expression in TNBC cells could provide an advantage during carcinogenesis, we silenced the expression of CD95 in human (MDA-MB-231) (Fig EV1A) and mouse (4T1) (Qadir *et al*, 2021) TNBC cells. Interestingly, the loss of CD95 in human (Fig 1A, Table 1) or mouse (Fig 1B, Table 2) TNBC cells induced a transcriptomic signature revealing 148 genes deregulated between parent and CD95-KO TNBC cells with a fold-change cutoff of 1.5 (Fig 1C and Table 3). These 148 genes were annotated for INFα, INFγ and TNFα pro-inflammatory responses determined by Gene Set Enrichment Analysis (GSEA) (see Tables 1, 2 and 3).

**Figure 1.**
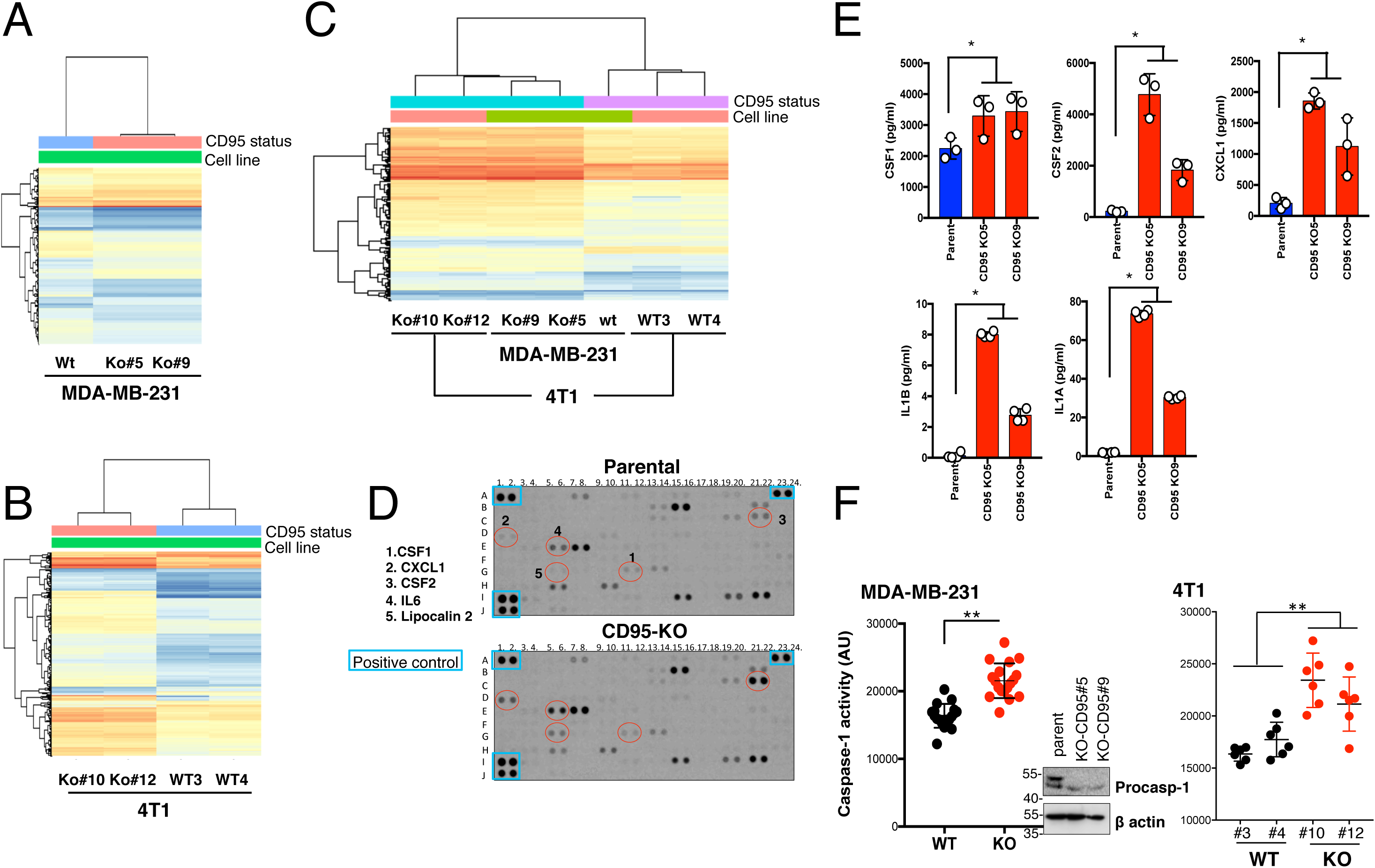
Loss of CD95 induces an inflammatory transcriptomic signature in TNBC cells. **A.** Heatmap of the genes differently expressed (386 genes with FC ≥1.5 or FC≤-1.5 and p.value≤0.05) between parental TNBC MDA-MB-231 cell line and two CD95-KO counterparts. **B.** Heatmap of the genes differently expressed (244 genes with FC ≥1.5 or FC≤-1.5 and p.value≤0.05) between two parental TNBC 4T1 clones and two CD95-KO counterparts. **C.** Heatmaps of the 148 genes modulated in a similar fashion in mouse (4T1 cells) and human (MDA-MB-231 cells) TNBC cells and their respective CD95 KO cells. **D.** The expression level of 102 different cytokine was evaluated using Proteome Profiler™ Human XL Cytokine Array Kit in supernatants of parental TNBC cells and their CD95-KO counterpart. Image is representative of three independently performed experiments. **E.** Certain cytokines up-regulated in the Proteome Profiler™ Human XL Cytokine Array Kit in D. were dosed by ELISA in supernatants of CD95-KO TNBC cells as compared to those of parental counterpart. Mean ± SD, p-values were calculated using Nonparametric Mann-Whitney test (n=3). **F.** The caspase-1 activity was measured in indicated MDA-MB-231 cells (*left panel,* n=18) or 4T1 cells (*right panel,* n=6). Mean ± SD, p-values were calculated using Nonparametric Mann-Whitney test. *Inset:* Parental MDA-MB-231 and their CD95-KO counterparts were lysed and indicated immunoblotting were performed. β-actin served as a loading control.

**Table 1.** Gene differently expressed between CD95-ko and wild type MDA-MB-231 cells.

**Table 2.** Gene differently expressed between CD95-ko and wild type 4T1 cells.

**Table 3.** Gene differently expressed between CD95-ko and wild type TNBC cells (4T1 and MDA-MB-231 cells)

To validate the inflammatory signature associated with CD95 loss in TNBC cells, we next conducted a holistic analysis of cytokines whose expression changed in CD95-KO MDA-MB-231 cells as compared to their wild type counterpart (Fig 1D). Among the 105 secreted factors, a Human proteome profiler cytokine array revealed that 28 cytokines were over-expressed with a cut-off greater than or equal to 2 and 12 cytokines down-regulated more than or equal to 2-fold (Fig 1D and Table 4). ELISAs confirmed the secretion of inflammatory cytokines including IL1α, IL1β, CSF1, CSF2 and CXCL1 in supernatants of CD95-KO cells as compared to those of their wild type counterpart (Fig 1E). Of note, IL-1β and IL-18 are produced as precursors requiring a caspase-1-driven proteolytic cleavage by the inflammasome to be secreted as active cytokines (Martinon *et al*, 2002). Because IL-1β was secreted when CD95 was deleted in MDA-MB-231 cells (Table 1), we evaluated whether CD95 loss could induce inflammasome activation in TNBC cells to produce the active and secreted forms of IL-1β and IL-18. Caspase-1 activity was increased in human and mouse CD95-KO TNBC cells as compared to their parental counterparts (Fig 1F) indicating that the loss of CD95 in TNBC cells caused a two-step pro-inflammatory response involved in activation of the inflammasome and secretion of cytokines.

**Table 4.** Cytokine expression in CD95-ko and wild type TNBC cells. The expression level of the indicated cytokines was quantified using Proteome Profiler Human XL Cytokine array (see *Materials and Methods* session). For each experiment, membrane dots have been quantified by densitometry to calculate mean and SD of three experiments performed independently.

Although we failed to detect m-CD95L at the plasma membrane of human (MDA-MB-231, SUM159) and mouse (4T1) TNBC cells (Fig EV1B), we could not rule out that an undetectable amount of m-CD95L still engaged CD95 in an autocrine fashion to induce a basal signaling pathway, which was abrogated by the loss of CD95 in cancer cells. To evaluate this possibility, we knocked-out CD95L in 4T1 TNBC cells. The selected cells that harbored an inserted base pair (adenine) within the second CD95L codon leading to a frameshift and the synthesis of a truncated peptide (Fig EV1C). No significant difference in the CSF1, CSF2, CXCL1, IL1α and IL1β transcription was measured between parental and two CD95L-KO 4T1 cells (Fig EV1D) indicating that elimination of the ligand did not recapitulate the inflammatory phenotype observed with CD95 loss. In addition, we incubated mouse 4T1 cells with the neutralizing anti-CD95L mAb MLF4 and human MDA-MB-231 cells with the antagonist anti-CD95L mAb NOK1, respectively. Interestingly, the 5-gene pro-inflammatory signature remained unaffected when TNBC cells were incubated for 24 hours in the presence of neutralizing anti-CD95L mAbs MFL4 or NOK1 (Fig EV1E). Overall, these findings indicated that elimination of CD95 in TNBC cells unleashed a pro-inflammatory signaling pathway through a ligand-independent mechanism.

### A NF-κB-dependent signaling pathway is activated in TNBC cells devoid of CD95

The differential transcriptomic signature obtained by comparing wild type with CD95-KO TNBC cells was strongly related to NF-κB activation including TNF-α signaling via NF-κB in gene-set enrichment analysis (GSEA) (Tables 1, 2 and 3). In agreement with this, elimination of CD95 in MDA-MB-231 cells led to a potent activation of the NF-κB signaling pathway as monitored by the phosphorylation of IKKα/β at Ser176/177 and Ser180/181, IκBα at Ser32, and p65 at Ser536 (Sakurai *et al*, 1999) (all hallmarks of NF-κB activation) (Fig 2A). The five members of the NF-κB family including RelA (p65), RelB, c-Rel, NFκB1 (p50/p105), and NFκB2 (p52/p100) share a conserved Rel homology domain (RHD) responsible for DNA binding. However, only p65, RelB, and c-Rel contain a transactivation domain in their C-terminal regions. Because, p50 is devoid of transactivation domains, nucleus accumulation of p50/p50 homodimers is considered as a transcriptional repressor of the NF-κB response (Grundstrom *et al*, 2004; Kang *et al*, 1992; Kravtsova-Ivantsiv *et al*, 2015; Udalova *et al*, 2000; Zhong *et al*, 2002). Although the quantity of p50 was slightly increased in nuclei of CD95-KO cells when compared to parental MDA-MB-231 cells (Fig 2B-C), this was less pronounced when compared to nuclear accumulation of p65 (Fig 2B-C). In agreement with the accumulation of an active p50/p65 heterodimer in nuclei of CD95-KO TNBC cells, an NF-κB reporter confirmed that the loss of CD95 in human (MDA-MB-231 and SUM159) and mouse (4T1) TNBC cells was associated with the induction of this signaling pathway (Fig 2D).

**Figure 2.**
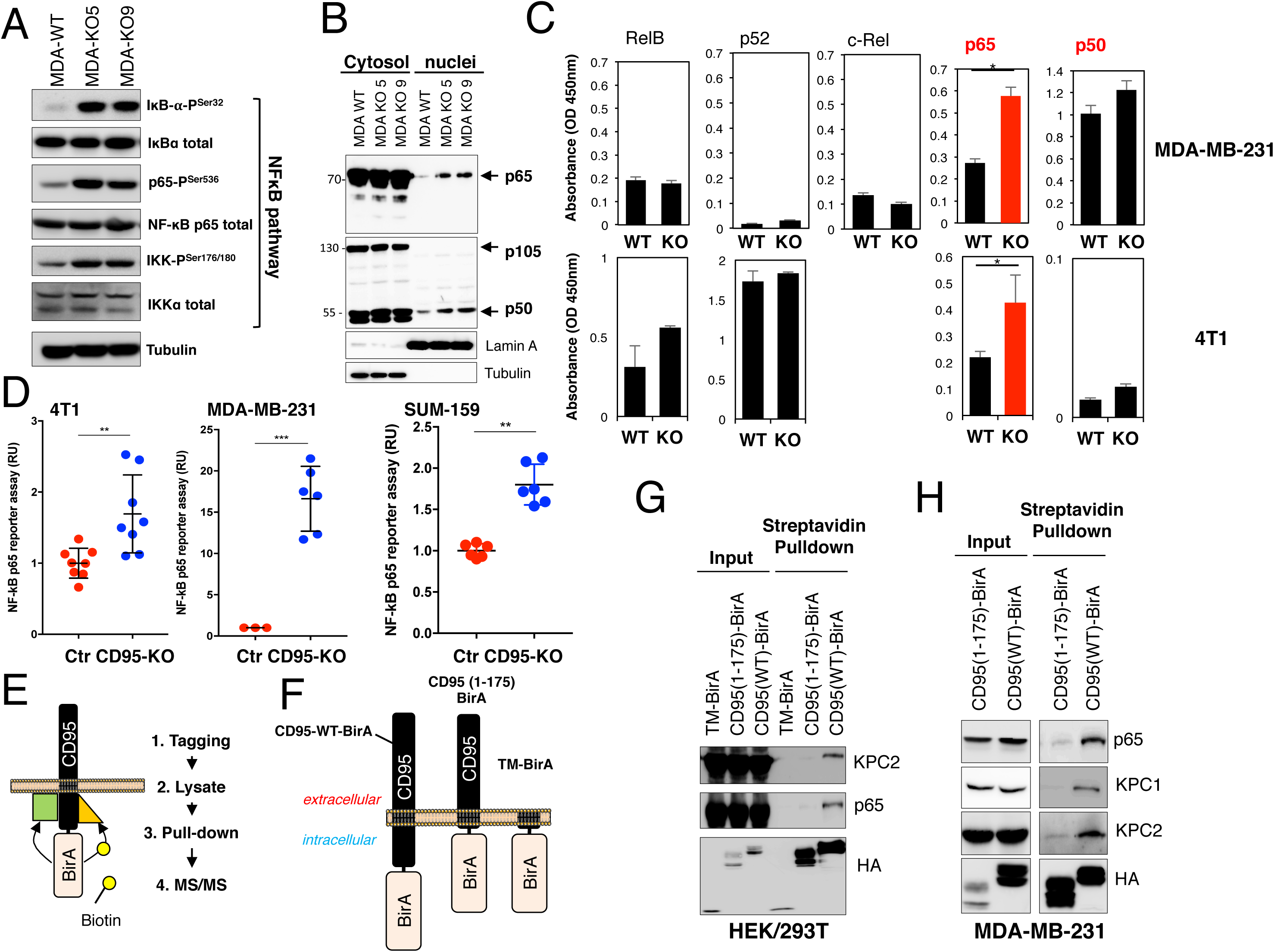
KPC2 and p65, two novel interactants of CD95. **A.** The activation status of the NF-kB signaling pathway was analyzed in wild-type (WT) and CD95 knock-out (KO5 and KO9) MDA-MB-231 cells by immunoblotting with the indicated antibodies. Tubulin immunoblot serves as a loading control. **B.** The presence of p65, p105 and p50 in the whole lysate (cytosol) or the nucleus fraction of wild type and CD95-KO cells was evaluated by immunoblotting. Lamin A and tubulin serve as loading control for nucleus and cytosolic fractions, respectively. **C.** Nuclear extracts from WT and CD95-KO cells were subjected to the indicated ELISA to quantify activation of NF-kB. Mouse c-Rel DNA binding cannot be assessed with this kit. Mean ± SEM (n=3), p-values were calculated using Nonparametric Mann-Whitney test. **D.** RelA activity was measured in indicated tumor cells using luciferase reporter assay (n=6-8)ù. ** and *** stand for p<0.001 and p<0.0001, respectively using unpaired and non-parametric Mann-Whitney t-test. **E.** Schematic representation of the BioID experiment. **F.** Schematic representation of BirA-fused CD95 constructs. **G-H.** BioID assay was conducted in CD95-KO HEK293T cells (G) and MDA-MB-231 cells (H) and after streptavidin-pull-down indicated immunoblotting were performed. Images are representative of three independent experiments.

These findings raised the question of how CD95 expression could control the NF-κB response and how its loss in TNBC cells triggered this inflammatory signaling pathway. To address this, we conducted a comprehensive proteomic analysis of the CD95 interactome using the proximity-dependent Biotin identification (BioID) method (Roux *et al*, 2012). Briefly, the activated E. coli BirA biotin ligase (mutant R118G) was fused to CD95 (Fig 2E). This enzyme has the ability to biotinylate the surrounding proteins located within a radius of 10 nm. Three constructs consisting of wild type CD95, a CD95 devoid of its intracellular region (CD95(1-175)-BirA) and the transmembrane domain of CD95 (TM-BirA) were fused to BirA (Fig 2F) and transiently expressed in CD95-KO HEK293T cells. By fusing CD95 to this BirA domain, even proteins interacting weakly and transiently with the receptor can be biotinylated in an irreversible fashion and thereby detected by mass spectrometry analysis (MS). Cells were lysed and biotin-conjugated proteins were next precipitated using streptavidin beads and identified by MS/MS. Among the 198 proteins selectively tagged by BirA and associated with the wild type CD95 construct, identification of caspase-8 and CD95 validated the method to characterize the CD95 interactome (Table 5). Of note, some biotinylated factors corresponded to inhibitors of the NF-κB pathway including COMMD7 (Esposito *et al*, 2016), KPC2 (Kravtsova-Ivantsiv *et al*., 2015) or TRAF-D1 (Sanada *et al*, 2008) (Table 5). Also, the transcription factor p65 was detected (Table 5). Pull-down experiments confirmed that KPC2 and p65 were associated with wild type CD95 and these interactions were lost with CD95(1-175) devoid of its intracellular region (Fig 2G). Reconstitution of CD95-ko MDA-MB-231 cells with CD95-BirA also revealed the recruitment of KPC2 and KPC1 within a CD95 complex (Fig 2H). On the other hand, TRAF-D1 and COMMD7 were not detected in HEK293T and MDA-MB-231 cells indicating that either these factors were present in CD95 complex but to a lesser extent as compared to KPC2 and p65 or that they corresponded to false positive hits. NFκB1 (p105) is synthesized as a large precursor, which is processed to generate the NF-κB subunit p50. The KPC2/KPC1 complex is involved in p105 ubiquitination to promote its partial degradation and the accumulation of its N-terminal product, namely p50, forming homodimers, which inhibit the inflammatory functions of the heterodimer p50/p65 (Kravtsova-Ivantsiv *et al*., 2015).

**Table 5.** Proteomic analysis of BioID. Biotin-labelled proteins differently pulled-down in HEK cells transfected with WT (CD95(WT)-BirA) or Mut2 corresponding to either a CD95-BirA construct devoid of its intracellular region (CD95(1-175)-BirA) or the CD95 transmembrane domain fused to BirA (TM-BirA))

To identify the CD95 domain involved in the recruitment of KPC2 and p65, we next conducted co-immunoprecipitation experiments in HEK293T cells. Although immunoprecipitation of wild type CD95 confirmed its association with both KPC2 and p65 (Fig 3A and 3B), elimination of the C-terminal region of CD95 led to a loss of these interactions suggesting that p65 and KPC2 were recruited via the C-terminal region of CD95 encompassing amino acid residues 303 to 319 (Fig 3A and 3B). Furthermore, only the amino terminal region of p65 (amino acid residues 1 to 307) encompassing its nuclear localization sequence (NLS) interacted with CD95 (Fig 3B) suggesting that CD95 could prevent nuclear-localization of p65 by masking its NLS similarly to inhibitor of kappa B (IκB) (Beg *et al*, 1992). To confirm that p65 and KPC2 required the C-terminal region of CD95 to be recruited, we synthesized and purified the different CD95 domains including CID, DD and C-term fused to glutathione S-transferase (GST), and incubated them with lysates of HEK293T cells expressing KPC2 or p65. As shown in Figure 3C, only pull-down of the C-terminal region of CD95 allowed formation of a complex containing p65 and KPC2. In agreement with these experiments, immunoprecipitation of KPC2 validated the existence of a complex containing CD95 and p65 (Fig 3D) and similarly, p65 pull-down showed the presence of CD95 and KPC2 in the immune complex (Fig 3D). To determine whether KPC2 and p65 interactions with CD95 were direct, we next performed a GST pull-down assay. CD95-GST interacted with purified KPC2 but failed to pull-down p65 (Fig 3E). Interestingly, in the presence of KPC2, CD95 was able to recruit p65 (Fig 3E) suggesting that KPC2 could serve as an adapter molecule for the recruitment of the NF-κB factor. To confirm that the KPC2/CD95 interaction was direct, we next developed a protein-fragment complementation assay (PCA) (Stefan *et al*, 2007), in which the Renilla luciferase enzyme was split into two fragments (F1 and F2) and fused to different proteins. When the proteins interact, their interaction favors luciferase refolding and the subsequent recovery of its enzymatic activity (Fig 3F). This method confirmed the direct interaction between KPC2 and CD95 (Fig 3F). In agreement with the pull-down experiments, PCA also indicated that both the intracellular region and the C-terminal region of CD95 bound full length KPC2, while DD or CID-CD95 failed to do it and thereby to refold the Renilla luciferase and allow light emission (Fig EV2A). A PCA competitive assay was developed (Fig EV2B) and established that the CD95/KPC2 interaction could only be disrupted by the over-expression of the intracellular region of CD95 or its C-terminal region (303-319 residues) (Fig EV2B-C), indicating that KPC2 is a previously unrecognized binding partner of the CD95 C-terminal region.

**Figure 3.**
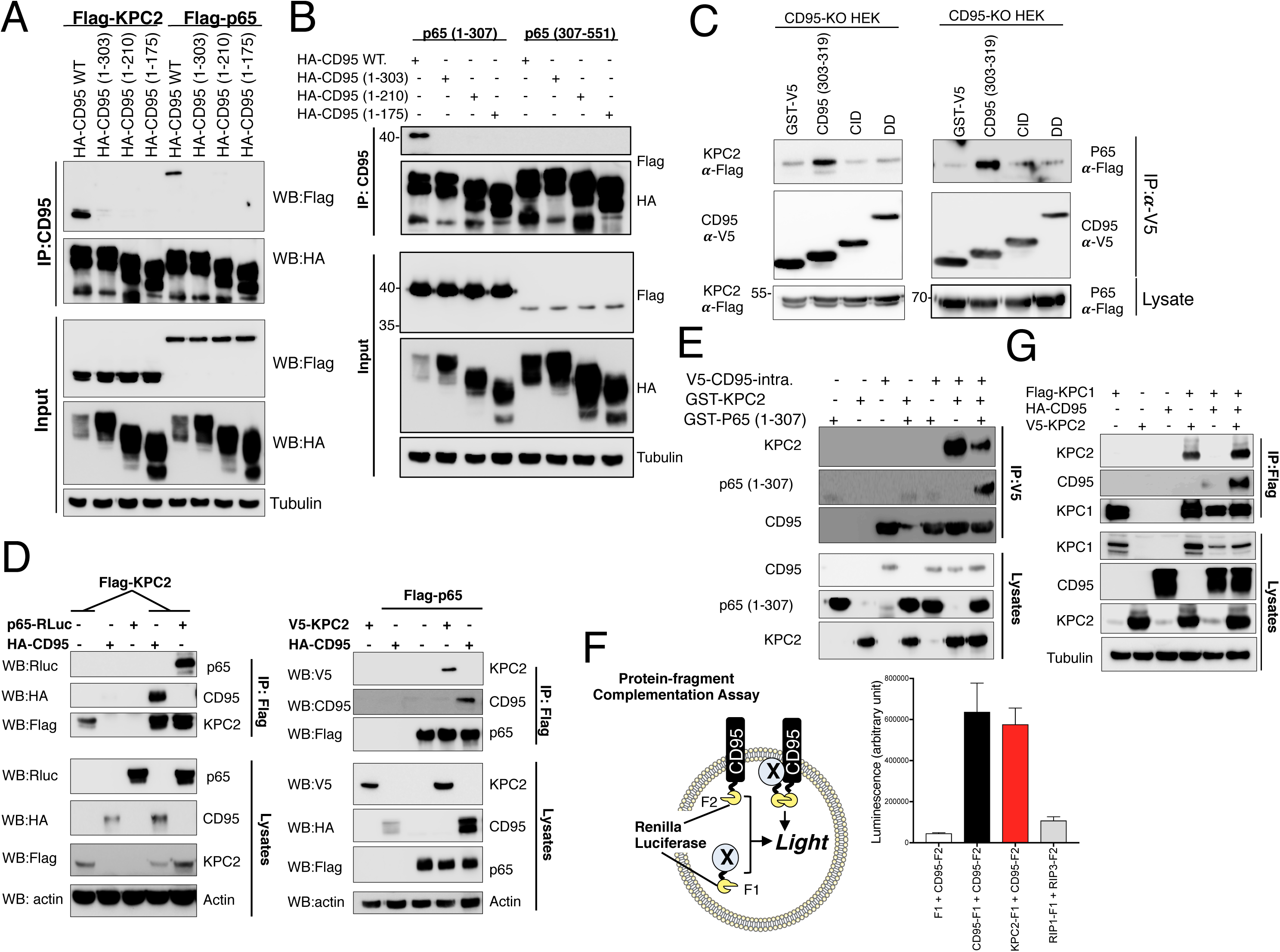
KPC2 binds the C-terminal region of CD95 and is an adaptor for p65 and KPC1. **A.** CD95-KO HEK293T cells were transfected with Flag-KPC2 or Flag-p65 and indicated HA-tagged CD95 constructs. After 24 hours, cells were lysed, HA immunoprecipitations were performed and the immune complex was resolved by SDS-PAGE. Indicated immunoblotting were performed. Images are representative of three independent experiments. **B.** CD95-KO HEK293T cells were co-transfected with Flag-tagged N-term (1-307) or C-terminal (307-551) p65 and with indicated HA-CD95 constructs. 24 hours after transfection, cells were lysed and HA immunoprecipitations were performed. Immune complex was resolved by SDS-PAGE and indicated immunoblotting were conducted. **C.** Indicated GST-V5-CD95 domains (100 ng) were produced in E.*coli*. and mixed with lysate (1 mg) of HEK293T cell transfected either Flag-KPC2 or Flag-p65. CD95 domains were immunoprecipitated using anti-V5 mAb and the presence of KPC2 or p65 was evaluated by immunoblot. Images are representative of three independent experiments. **D.** CD95-KO HEK293T cells were co-transfected with Flag-KPC2 (left) or Flag-p65 (right) and HA-CD95 WT in the presence of p65-Renilla luciferase (domain F1, left panel) or V5-KPC2 (right panel). After 24h, Flag immunoprecipitations was performed and the immune complex was resolved by SDS-PAGE. Indicated immunoblotting were performed. **E.** GST-KPC2, p65 (1-307) and different GST-V5 CD95 domains were produced in E.*coli* and purified. Indicated proteins (100 ng) were mixed for 1h and V5 immunoprecipitation was performed. The immune complex was resolved by SDS-PAGE and indicated western blots were performed. Images are representative of three independent experiments. **F.** Protein-fragment complementation assay (PCA). HEK293T cells were co-transfected with CD95 fused to the C-term region of Renilla luciferase (F2) and the N-term part of Renilla luciferase (F1) fused to indicated proteins and luciferase activity was measured. RIP1-F1 and RIP3-F2 interaction served as a positive control. Data represent mean ± SD (n=3). **G.** CD95-KO HEK293T cells were co-transfected with Flag-KPC1, HA-CD95 or V5-KPC2. After 24 hours, KPC1 was immunoprecipitated and the immune complex was resolved by SDS-PAGE and indicated western blots were realized. Images are representative of three independent experiments.

In the Kip1 ubiquitination-promoting complex (KPC) consisting of KPC1 and KPC2, KPC1 polyubiquitinates the precursor p105 leading to its partial degradation giving rise to the p50 subunit (Kravtsova-Ivantsiv *et al*., 2015). In HEK293T cells, KPC1 immunoprecipitation confirmed its association with KPC2 (Fig 3G). The immunoprecipitation of KPC1 also revealed a small quantity of CD95, probably due to the low endogenous level of KPC2 present in HEK293T cells (Fig 3G). Indeed, this amount dramatically increased when KPC2 was co-transfected (Fig 3G) suggesting that like p65, KPC1 was indirectly recruited by CD95 through its interaction with KPC2. Strikingly, CD95 deletion resulted in an accumulation of p105 when compared to parental cells (Fig 4A) without affecting the expression level of p65 (Fig 4A). In agreement with its KPC1 stabilizer role (Kravtsova-Ivantsiv & Ciechanover, 2015), KPC2 deletion in CD95-KO cells reduced the expression level of KPC1 (Fig 4A). Interestingly, the double knock-out of CD95 and KPC2 did not translate into a more pronounced accumulation of p105 when compared to CD95-KO cells (Fig 4A) suggesting that at least in tumor cells, the KPC1/KPC2-mediated regulation of p105 expression mainly relies on CD95 expression. We next wondered whether CD95 participated in p105 ubiquitination and co-transfected Flag-p105 with HA-tagged ubiquitin in HEK293T cells, after knockout of CD95 alone or of CD95 and KPC2 (Fig 4B). Immunoprecipitation of p105 revealed that although a small amount of the over-expressed proteins underwent ubiquitination, this process was dramatically reduced by the loss of CD95 (Fig 4B). Combination of KPC2 and CD95 deletion did not significantly enhance this reduction (Fig 4B) supporting that the presence of CD95 was required for KPC1/KPC2-driven ubiquitination of p105. In agreement with the CD95-KO data, elimination of KPC2 led to the secretion of pro-inflammatory cytokines (Fig EV2D).

**Figure 4.**
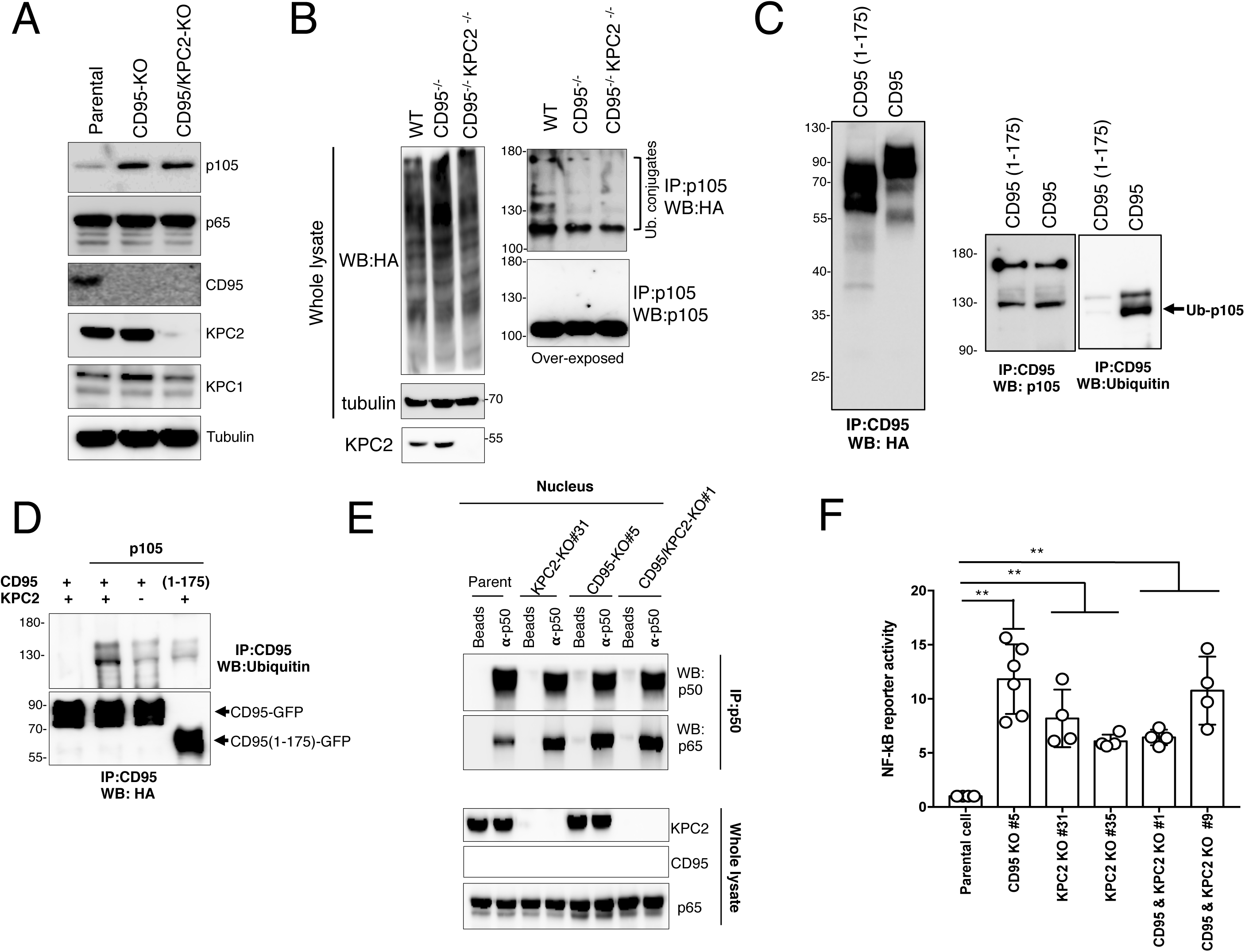
CD95 loss promotes the partial degradation of p105 and the nucleus accumulation of p50 homodimers. **A.** CD95 and KPC2 were knocked out in HEK293T cells using CRISPR/Cas9 and the expression level of the KPC1/KPC2 substrate p105 was analyzed by immunoblot. Images are representative of three independent experiments. **B.** CD95 and CD95/KPC2-KO HEK293T cells were co-transfected with cDNAs encoding for p105 and HA-Ubiquitin. After 24 hr, the proteasome inhibitor MG132 was added for 3 hr, cells were lysed and p105 was immunoprecipitated. Total lysate (left panels) or immunoprecipitates (right panels) are depicted. Following SDS-PAGE, indicated immunoblots were performed. Images are representative of three independent experiments. **C.** HEK293T cells were transfected with full length or intracellular-truncated (1-175) CD95, and CD95 complexes were immunoprecipitated and subjected to an *in vitro* ubiquitin conjugation assay in a reconstituted cell-free system in the presence of recombinant human p105 as described in *Materials and methods*. Then proteins were resolved by SDS-PAGE and indicated immunoblots were performed. Data are representative of three independent experiments. **D.** Parental HEK293T cells were transfected with full length or intracellular-truncated (1-175) CD95. KPC2-KO HEK293T cells were transfected with full length CD95. Then, cells were lysed and CD95 was immunoprecipitated. The immune complexes were subjected to an *in vitro* ubiquitin conjugation assay as depicted in C and next, proteins were resolved by SDS-PAGE and indicated immunoblots were performed. Data are representative of three independent experiments. **E.** p50 was immunoprecipitated in the nuclei of parental and CD95-KO HEK293T cells. The immune complex was resolved by SDS-PAGE and co-association with p65 was evaluated by immunoblotting. Data are representative of three independent experiments. **F.** RelA activity was measured in indicated tumor cells using luciferase reporter assay. ** stands for p<0.001 using unpaired and non-parametric Mann-Whitney t-test.

Finally, we monitored ubiquitin conjugation of wild-type p105 in a reconstituted cell-free system from immunoprecipitation of full length CD95 or CD95(1-175) constructs transfected in HEK293T cells. As shown in Figure 4C, p105 was efficiently ubiquitinated when incubated with the immune complex containing full-length CD95 (Fig 4C), while immunoprecipitation of a CD95 devoid of its intracellular region (CD95(1-175)) failed to add ubiquitin moieties to p105 (Fig 4C). Of note, although the same quantity of CD95 was immunoprecipitated in KPC2-KO HEK293T cells as compared to their parental counterpart (Fig 4D), the magnitude of p105 ubiquitination significantly dropped in the CD95-containing immune complex devoid of KPC2 (Fig 4D). Removal of the intracellular domain of CD95 also dramatically reduced p105 ubiquitination (Fig 4C-D), confirming that CD95/KPC2 interaction caused ubiquitination and partial degradation of p105 (Fig 4D).

Because p50 does not contain a transactivation domain (TAD) and its homodimeric form mostly acts as a transcriptional repressor (Christian *et al*, 2016), we next wondered whether the CD95-driven ubiquitination of p105 could favor the accumulation of p50 homodimers in the nucleus at the expense the transcriptional activator p50/p65. To tackle this question, we isolated nuclei from parental, KPC2-KO, CD95-KO and double KO MDA-MB-231 cells and immunoprecipitated p50 (Fig 4E). While the immunoprecipitation of p50 from nuclei of CD95-expressing cells revealed a faint quantity of co-associated p65 in parental cells, this quantity dramatically increased in nuclei of CD95, KPC2 and double KO cells, confirming that loss of both CD95 or KPC2 caused a shift from inactive p50 homodimers to the active NF-κB heterodimer p50/p65 (Fig 4E). Confirming the transcriptional activity of the p50/p65 heterodimer, elimination of KPC2 or CD95 in TNBC MDA-MB-231 cells unleashed an NF-κB response (Fig 4F) that was similar to that observed in CD95/KPC2-double KO cells, supporting the observation that these two factors share the same signaling pathway to modulate the NF-κB transcriptional activity (Fig 4F).

Overall, these findings demonstrate that KPC2 binding to CD95 not only contributes to the ubiquitination and partial degradation of p105 but also sequesters its transcriptionally active partner p65 to the plasma membrane. Deletion of CD95 therefore shifts the balance from canonical NF-κB activation to activation of a noncanonical NF-κB pathway that causes induction of inflammatory chemo- and cytokines. This activity does not appear to require CD95L.

Stimulation of CD95 on signaling competent but apoptosis resistant cancer cells results in the activation of multiple nonapoptotic signaling pathways that include NF-κB which drives tumor promotion (Legembre *et al*, 2004). We now show that the role of CD95 and its link to the NF-κB signaling pathway on signaling incompetent cancer cells appears to be different. Our study suggests that in unstimulated cancer cells, CD95 forms a complex harboring KPC2/KPC1 and p65 favoring the generation of p50-p50 homodimer at the expense of its tumorigenic p50-p65 couple. Although the p50 homodimer may act as a transcriptional repressor because it lacks a transactivation domain (May & Ghosh, 1997), it can also interact with different transcriptional modulators, such as Bcl-3 (Fujita *et al*, 1993), p300 (Deng & Wu, 2003), or HMGA1/2 (Perrella *et al*, 1999) involved in chromatin remodeling. Accordingly, an imbalance of the p50 expression may shift the composition of NF-κB dimers, resulting in different cellular response to environmental stresses.

Our study identifies a novel mechanism to explain the pro-oncogenic role of CD95 in apoptosis resistant, and CD95 signaling incompetent TNBC cells. It involves sequestering a fraction of p65 at the plasma membrane and simultaneously promotes partial degradation of p105 to form p50, favoring the formation of p50 homodimers at the expense of the pro-inflammatory p50/p65 complex (Kravtsova-Ivantsiv *et al*., 2015). Because the degree of tumor-infiltrating CD8+ T-cells is a good prognostic marker in TNBC patients (Ali *et al*, 2016; Ali *et al*, 2014) and CD95 expression is mainly expressed by tumor-infiltrating leukocytes including CD8+ T-cells, it is not surprising that CD95 expression levels have been recently associated with worse relapse free survival and overall survival in TNBC women (Blok *et al*., 2017). Combined with our previous study in TNBCs (Qadir *et al*., 2021), we highlight a complex picture in which although the CD95 loss by tumor cells in TNBC patients can curtail the anti-tumor effect induced by CD95L-expressing T-and NK cells, it also induces an inflammatory gene signature resulting in a more pronounced NK-mediated anti-tumor response (Qadir *et al*., 2021). The expression of pro-inflammatory chemokines including CSF1, CXCL1 and CXCL2, may also promote the recruitment of macrophages and the CD95 loss by TNBC cells also drives the over-expression of lipocalin 2 (Lpn2) (see Fig 1D and Table 4), a small protein known to promote differentiation of macrophages to an anti-tumor M1 transcriptomic signature (Cheng *et al*, 2015) and to enhance their phagocytic activity (Toyonaga *et al*, 2016). All these processes combined may contribute to the complex anti-tumor response in TNBC cells devoid of CD95 we recently described (Qadir *et al*., 2021).

A comprehensive genomic analyses of breast cancer (Cancer Genome Atlas, 2012) cells previously identified the transcription factors, FOXM1 and c-myc which are hyperactivated in TNBCs as factors to contribute to the down-regulation of the CD95 expression (Penke *et al*, 2018; Zhang *et al*, 2017). Furthermore, breast cancers in general up-regulate the PI3K signaling pathway through the expression of constitutive active or amplification of PIK3CA, deletion of PTEN or INPP4B or Akt mutations (Nik-Zainal *et al*, 2016). This oncogenic signaling pathway activates the transcription repressor YY1 shown to be involved in the down-regulation of CD95 mRNA expression (Garban & Bonavida, 2001). Interestingly, although these transcription regulations might explain the reduction of CD95 observed in many tumors, the estrogen receptor-negative TNBC cells maintain a higher level of CD95 as compared to other breast tumors (Blok *et al*., 2017). TNBC cells therefore may maintain high CD95 expression to prevent the induction of an NF-κB-driven pro-inflammatory program favoring their immune destruction.

In conclusion, our work adds a novel immunological role to the cancer relevant activities of CD95. While the classical tumor evasion hypothesis associated with the CD95 loss in tumor cells remains valid, this immune selection might be counteracted by the induction of a NF-κB-driven pro-inflammatory pathway, which could account for the activation of the NK cell-mediated anti-tumor responses (Qadir *et al*., 2021).

## Materials and Methods

### Cells lines

All cells were purchased from ATCC (Molsheim Cedex, France). The 4T1 cell line was cultured in RPMI supplemented with 8% heat-inactivated FCS (v/v) and 2 mM L-glutamine at 37°C/5% CO_2_. HEK293T, MDA-MB-231 and SUM-159 cells were cultured at 37°C/5% CO_2_ in DMEM supplemented with 8% heat-inactivated FCS and 2 mM L-glutamine.

### Reagents and antibodies

3-[4,5-dimethylthiazol-2-yl]-2,5-diphenyltetrazolium bromide (MTT) was purchased from Sigma-Aldrich (L’Isle-d’Abeau-Chesnes, France). Antibodies against phospho-IκBα Ser32 (#2859), IκBα (#4814), phospho-IKK Ser176/177 (#2078), IKKα(#11930), phospho-p65 Ser536 (#3033), p65 (#8242), p105/p50 (#12540), p27 (#3686) were purchased from Cell Signaling Technology. Antibodies against Lamin A (sc-20680) and KPC1 (sc-101122) were from Santa Cruz. KPC2 antibody was purchased from Abcam (ab177619) and antibodies against Flag (clone M2) and α-Tubulin (clone DM1A) were from Sigma Aldrich. Anti-HA.11 antibody was purchased from Biolegend and antibodies against V5 and GST tags were from ThermoFisher.

### Plasmids

For all CD95 constructs, the numbering takes into consideration the subtraction of the 16 amino-acid signal peptide sequence. The vector pcDNA3.1-PS-HA-hCD95(1-319) encodes the CD95 signal peptide followed by the human influenza hemagglutinin (HA) tag in frame with CD95 full length. Plasmids encoding the different CD95 mutants were previously described (Poissonnier *et al*., 2016). For all the constructs except the CD95(303-319) plasmids, we amplified the ORF genes by PCR and cloned them using the Gibson Assembly protocol (NEB). For the CD95 (303-319) plasmids, we directly ligated duplexed primers into the plasmid. The primers used for PCR are listed below. V5-tagged GST was generated by inserting between BamHI and EcoRI sites of pGEX6P1 the annealed V5 tag (GKPIPNPLLGLDST) using primers: GGTAAGCCTATCCCTAACCCTCTCCTCGGTCTCGATTCTACG and CGTAGAATCGAGACCGAGGAGAGGGTTAGGGATAGGCTTACC. For BioID experiments, wildtype or truncated CD95 sequence was extracted either from 5UTR-SP-HA-CD95 wt (1-319)-eGFP peGFP-N1 or 5’UTR-SP-HA-CD95 wt (1-175)-eGFP peGFP-N1 by NdeI and SmaI and inserted into the NdeI and AfeI sites of pMCS-BioID2-HA (Addgene #74224). The plasmid encoding the CD95 transmembrane domain fused in frame to the BirA was created by inserting the following annealed primers into the EcoRI and BamHI site of pMCS-BioID2-HA: AATTCTTGGGGTGGCTTTGTCTTCTTCTTTTGCCAATTCCACTAATTGTTTGGGTGG and GATCCCACCCAAACAATTAGTGGAATTGGCAAAAGAAGAAGACAAAGCCACCCC AAG. All primers were purchased from Eurogentec (Liège, Belgium).

The following sgRNA sequences were cloned within PX459-V2 plasmid and then transfected with lipofectamine 3000 in HEK293T, 4T1, SUM-159 and MDA-MB231 cells to generate cell lines deficient for CD95 or CD95L according to manufacturer’s instructions. sgRNA sequences target human CD95, 5’CACCGAGGGCTCACCAGAGGTAGGA3’ (Fwd), 5’AAACTCCTACCTCTGGTGAGCCCTC3’ (Rev); mouse CD95 5’CACCGCTGCAGACATGCTGTGGATC3’ (Fwd), 5’AAACGATCCACAGCATGTCTGCAGC3’ (Rev); mouse CD95L 5’CACCGGTAATTCATGGGCTGCTGCA3’ (Fwd), 5’AAACTGCAGCAGCCCATGAATTACC 3’ (Rev). After transfection and puromycin selection for 48h, genome-edited cells were cloned by limited dilution. To generate KPC2 or KPC1 knockout cells, sgRNA (XXXXXXXX for KPC1 or CGGCGGCGGGATGTTCGTGC for KPC2) were synthetized in vitro using EnGen sgRNA Synthesis kit (NEB) and purified using RNA clean and concentrator (Zymo Research). Cells were then transfected with sgRNA and the EnGen Cas9 NLS protein (NEB) using Lipofectamine RNAiMAX and according to manufacturer’s instructions. After 48h, cells were cloned by limiting dilution.

### Chemokine quantification

Cells (5.10^5^) were plated in 6-well plate and incubated for 4 days at 37°C. Media were then collected, filtered on 0.22 µm and immediately frozen. Cytokines level were measured using either the Proteome Profiler Human XL Cytokine array (R&D Systems) or Quantikine ELISA kits for dedicated cytokines (R&D Systems), according to manufacturer’s instructions.

### Q-PCR

For qPCR analysis of cytokines expression, total RNA was isolated from cell lines using NucleoSpin RNA Kit (Macherey-Nagel) and subjected to Real-Time PCR using the high-capacity cDNA reverse transcription kit (Applied Biosystems, ThermoFisher Scientific). qPCR was performed using SYBR Green PCR Master Mix (Applied Biosystems). Results reported as fold change was calculated using the Δct method relative to the housekeeping gene GAPDH for Human or HPRT for mouse. Next, ΔΔct were assessed by comparing wildtype and KO cells. Primers purchased from Eurogentec (Angers, France) are listed in Table n°1.

### Caspase-1 activity

Cells (50,000) were seeded in 96-wells white plate and caspase-1 activities were assessed using Caspase-Glo 1 Inflammasome Assay (Promega #G9951) according to manufacturer’s instructions.

### Ubiquitination

HEK293T cells were transfected with plasmids encoding for Flag-p105 or Flag-p27 together with the plasmid HA-Ubiquitin (Addgene #18712). After 24h, cells were treated for 3h in the presence of 20 µM of proteasome inhibitor MG132, washed twice with PBS and lysed in IP buffer (50 mM Tris pH 7.4, 150 mM NaCl, 2 mM EDTA, 1% Triton X-100 and proteases inhibitor) supplemented with 5 mM of freshly added N-Ethylmaleimide (NEM). After sonication, lysates were clarified and Flag-tagged protein were precipitated using anti-Flag M2 magnetic beads (Sigma Aldrich). After several washes in IP buffer, precipitated proteins were resolved by SDS-PAGE and immunoblotting was performed with the indicated antibodies.

### NF-κB ELISA

To quantify the level of NF-kB transcription factor activity, fractionation of MDA-MB-231 and 4T1 cells was first performed using the Nuclear Extract kit (Active Motif). The level of protein binding to DNA consensus sequence was then assessed by ELISA using the TransAM NF-kB Family kit (Active Motif) according to manufacturer’s instructions.

### Subcellular fractionation

Cell lysis and immunoblot analysis were performed as described previously (Poissonnier *et al*, 2018). Proteins were visualized by enhanced chemiluminescence using ECL revelblot (Ozyme) and scanned with LAS-4000 imager (Fujifilm). Subcellular protein fractionation was performed using the REAP protocol as described (Suzuki *et al*, 2010). Briefly, cells were washed in PBS, and resuspended in lysis buffer (PBS, 0.1% NP-40). Lysate was centrifuged for 10 sec at 13,000 rpm and the supernatant was collected (cytoplasmic fraction). Pellet was then washed in lysis buffer, centrifuged for 10 sec at 13,000 rpm and the pellet was resuspended in lysis buffer and sonicated (nuclear fraction).

### Protein production and Pulldown assay

GST-V5 encoding wildtype and truncated CD95, GST-KPC2 and GST-p65 (1-307) constructs were transformed in BL21 *E.coli* strain. Bacteria were grown at 37 °C in Luria Broth to an optical density of 0.8, and protein expression was induced with 0.5 mM isopropyl b-D-1-thiogalactopyranoside (IPTG) at 30°C for 4 hours. After centrifuging the cells, the pellets were resuspended in phosphate buffered saline (PBS) (137 mM NaCl, 2.7 mM KCl, 10 mM Na2HPO4, 2 mM KH2PO4, pH 7.4), containing 1 mM phenylmethylsulfonyl fluoride (PMSF), and 0.1 mg/ml lysozyme, and cells were lysed by sonication. Cell debris was removed by centrifugation, and the GST-fusion protein was purified using Glutathione-Sepharose affinity columns (GE Healthcare). The proteins were finally eluted in PBS containing 20 mM glutathione (pH 8) and the concentration and purity assessed by SDS-PAGE using Coomassie Brilliant Blue.

100 ng of CD95 constructs (GST-V5-CD95 (175-319), GST-V5-CD95 (175-210), GST-V5-CD95 (211-303) and GST-V5-CD95 (303-319)) and control GST-V5 were mixed with 100 ng of GST-KPC2 or GST-p65 (1-307) in IP Buffer (25mM Hepes, 150 mM NaCl, 2mM EDTA and 1% Triton X100 and protease inhibitors) for 1h at room temperature. Anti-V5-agarose beads (Sigma Aldrich) were then added and incubated for 2h at 4°C on rotating wheel. After 4x washes in IP buffer, the pulled-down complex was resolved by SDS-PAGE and analyzed by immunoblotting.

### Immunoprecipitation

For classical immunoprecipitations, indicated cells were washed once in phosphate-buffered saline and lysed in IP buffer. Protein lysates were incubated overnight with 5 µL of Flag or V5 antibody-conjugated beads (Sigma-Aldrich) or with 1 µl of p50 antibody together with 20 µl of Protein A magnetic beads (Life Technologies). After 4 washes in IP buffer, the beads were resuspended in 2X Laemmli’s buffer and subjected to immunoblotting analysis.

For immunoprecipitation of CD95 complexes, HEK293T cells were transfected with Flag-KPC2 or Flag-p65. After 24 h, transfected cells were lysed in Hepes Buffer (Hepes 25mM, NaCl 150mM, NaF 2mM, NaVO_4_ 1mM, EGTA 2mM). Cell lysates were incubated for 2h at 4°C with 100 ng of the different GST-V5-CD95 constructs and then overnight at 4°C with 10 µl of Anti-V5 Agarose Beads (Sigma-Aldrich). After extensive washing in Hepes buffer, the precipitated complex was resolved by SDS-PAGE and immunoblotting was performed with the indicated antibodies.

### BioId and Proteomic analysis

HEK293T cells were transfected with plasmids encoding either wild-type (CD95(WT)-BirA), CD95 devoid of its intracellular region (CD95(1-175)-BirA) or CD95 transmembrane domain (TM-BirA) fused in frame with BirA-R118G sequence. After 24h, Biotin (50 µM) was incubated with cells for 16 hours. After 3 PBS washes, cells were scraped in lysis buffer (50mM Tris pH 7.4, 500 mM NacL, 0,4% SDS, 1mM DTT, 2% Triton X-100 and protease inhibitors), sonicated. Cell lysate was incubated with High-Capacity Streptavidin Agarose beads (Pierce) overnight at 4°C. Beads were washed once in 2% SDS, twice in modified RIPA buffer (50 mM Tris pH7.4, 150 mM NaCl, 1mM EDTA, 0.1% SDS, 1% NP40), twice in TAP buffer (50 mM Hepes pH8, 150 mM NaCl, 2mM EDTA, 0.1% NP40, 10% glycerol) and once in PBS. Precipitated proteins were then analyzed by immunoblotting or mass spectrometry.

### NF-κB reporter assay

Indicated cell lines were transfected using lipofectamine 3000 with 3 µg of p65 reporter plasmid (pHAGE NF-κB-TA-LUC-UBC-GFP-W plasmid from Addgene #49343). After 48h, cells were lysed using 150 µl of Luciferase cell culture lysis buffer (Promega) and luminescence was assessed using 96-wells white plate. Luciferase activities were directly measured using Luciferase assay system (Promega) on TECAN infinite 200 PRO plate reader. GFP fluorescence, which is constitutively expressed by the reporter vector served to normalize the luminescent signal according to the transfection efficiency, and the p65 activity signals was depicted as luminescent intensity (reporter assay) / Fluorescent signal (GFP).

### Protein-fragment Complementation Assay (PCA)

HEK293T cells were electroporated with 10 μg of DNA using BTM-830 electroporation generator (BTX Instrument Division, Harvard Apparatus). 24 h after transfection, cells (10^6^) were washed and resuspended in 100 μl PBS placed in OptiPlate-96 (PerkinElmer, Waltham, MA, USA) in the presence or absence of indicated peptides (50 μM) for 1 hour. Coelenterazine-h (5 µM, Sigma-Aldrich) was then injected and the Renilla luciferase activity was assessed for the first 10 seconds using Infinite200Pro (Tecan, Männedorf, Switzerland).

### Microarray analysis

RNA quality was assessed using an RNA6000 nano chip (Agilent). For each condition, 9 ng of RNA were reverse transcribed using the Ovation PicoSL WTA System V2 (Nugen, Leek, The Netherlands). MDA-MB-231 fragmented cDNAs were hybridized to GeneChip Human Gene 2.0 ST microarrays (Affymetrix). 4T1 fragmented cDNAs were hybridized to Clariom S mouse microarrays (Affymetrix). Hybridized microarrays were scanned by a GeneChip Scanner 3000 7G (Affymetrix). Raw data and quality-control metrics were generated using Expression Console software (Affymetrix). Probes were normalized by robust multi-array averaging with rma function from oligo R package (Carvalho & Irizarry, 2010) and mapped using hugene20sttranscriptcluster.db and clariomsmousetranscriptcluster.db (MacDonald JW, 2017) as database for MDA-MB-231 and 4T1 datasets, respectively. For the parent and CD95-KO TNBC cells differential expression analysis, the two normalized datasets were merged by upper case gene symbol and a batch correction were made by ComBat function from Surrogate Variable Analysis R package (Leek *et al*, 2012). Raw and normalized data are deposited to the GEO database under accession ID GSEXXXXX and GSEXXXXX. Statistical analyses were performed using the Linear Model for Series of Arrays (lmFit) from limma R package (Ritchie *et al*, 2015); For analysis of 4T1 and MDA-MB-231 separately genes showing a minimum of 1.5 fold change in expression and a minimum P value of 0.05 were considered significant. For merged datasets analysis genes showing a minimum of 1.5 fold change in expression and a minimum P value of 0.1 were considered significant. Gene expression heatmaps were made with pheatmap R package.

### GSEA

Gene Set Enrichment analysis was performed by GSEA function from ClusterProfiler R (Yu *et al*, 2012) package with Homo sapiens hallmark gene sets data previously collected in msigdbr R package, gene symbol translated into entrezid using bitr function with org.Hs.eg.db and pvalue cut off was considered significant at 0.25.

### *In vitro* ubiquitin conjugation Assays

HEK293T cells were transfected in the presence of full length or truncated (1-175) CD95 constructs. After 24h, cells were lysed in TNET buffer (50 mM Tris pH7.4, 100 mM NaCl, 5 mM EDTA, 0,5% Triton X-100, protease inhibitors) and CD95 was immunoprecipitated using the anti-CD95 mAb APO1-3. After washes in TNET buffer, beads were incubated in the presence of recombinant human p105 (150 ng) (Novus Biologicals) for 1 h at 37°C with 50 µl of ubiquitination reaction buffer [5 mM Mg-ATP solution, ubiquitin activating enzyme E1 (100 nM, Enzo Life Sciences, Villeurbanne, France), E2 UbcH5c (2.5 µM, Enzo Life Sciences), biotinylated Ubiquitin (2.5 µM, Enzo Life Sciences), 2U inorganic pyrophosphatase (Sigma Aldrich), 1mM DTT]. The ubiquitination reaction was then stopped by the addition of 5X Laemmli buffer and samples were resolved by SDS-PAGE.

### Cloning strategy – vectors and primers

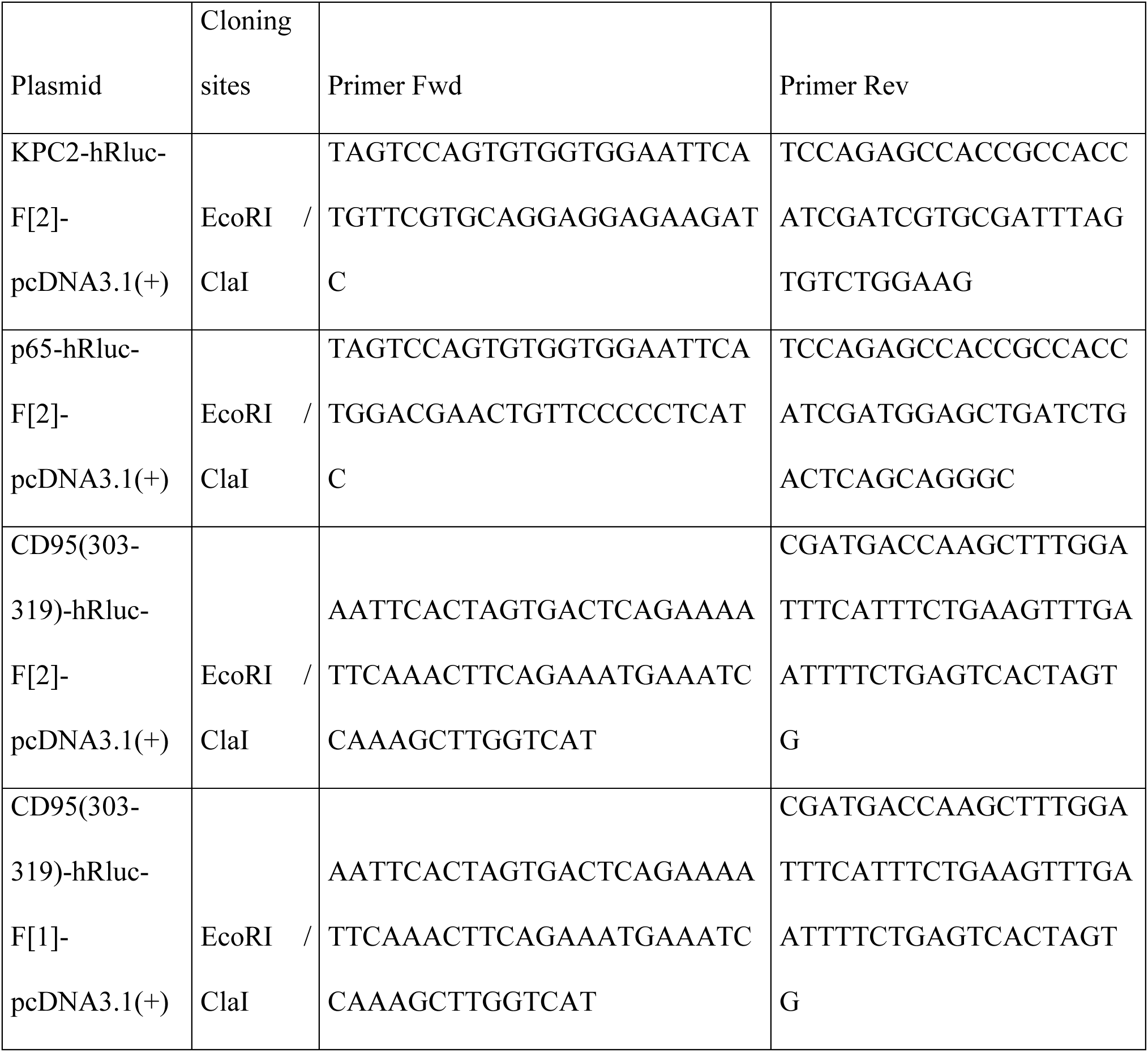
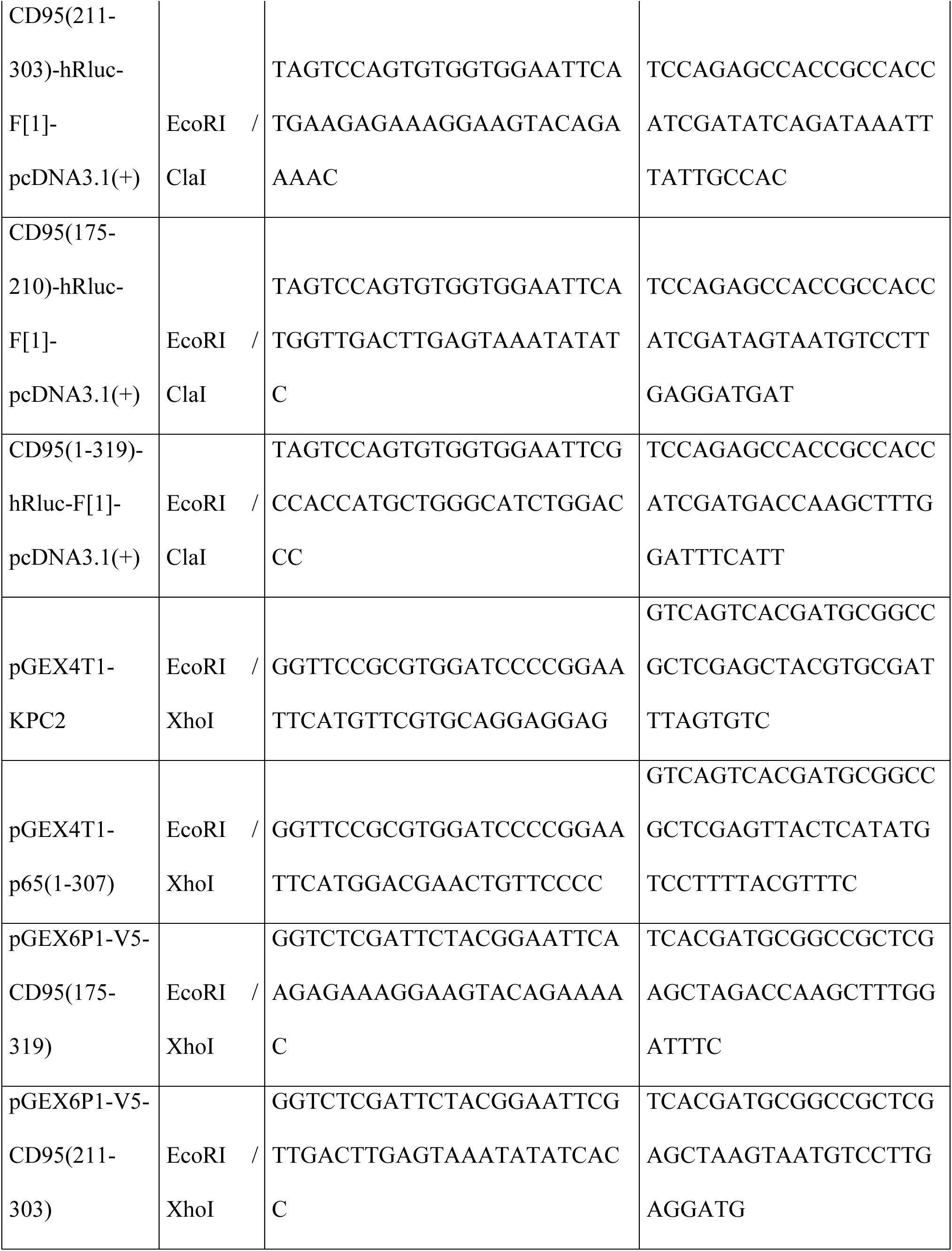
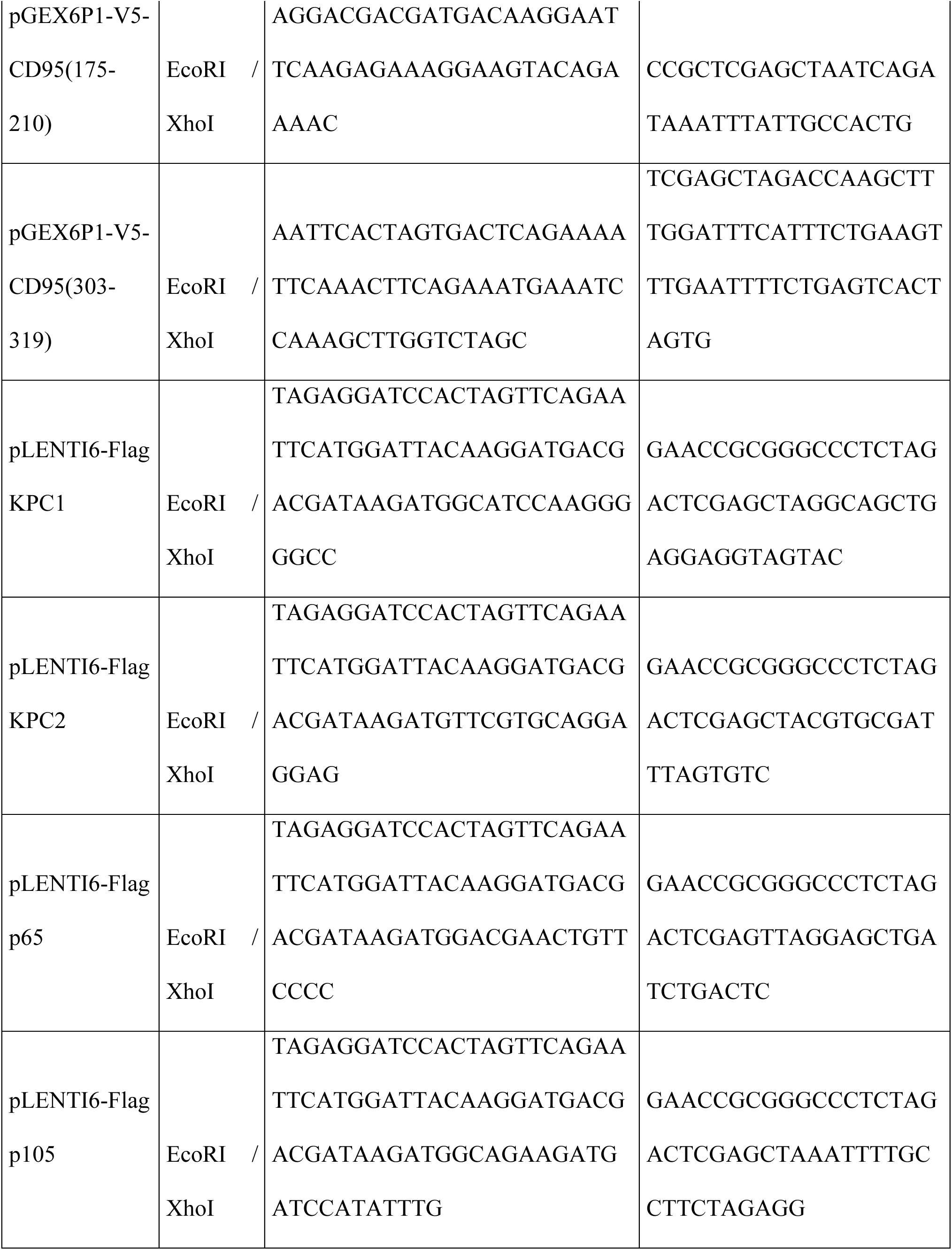
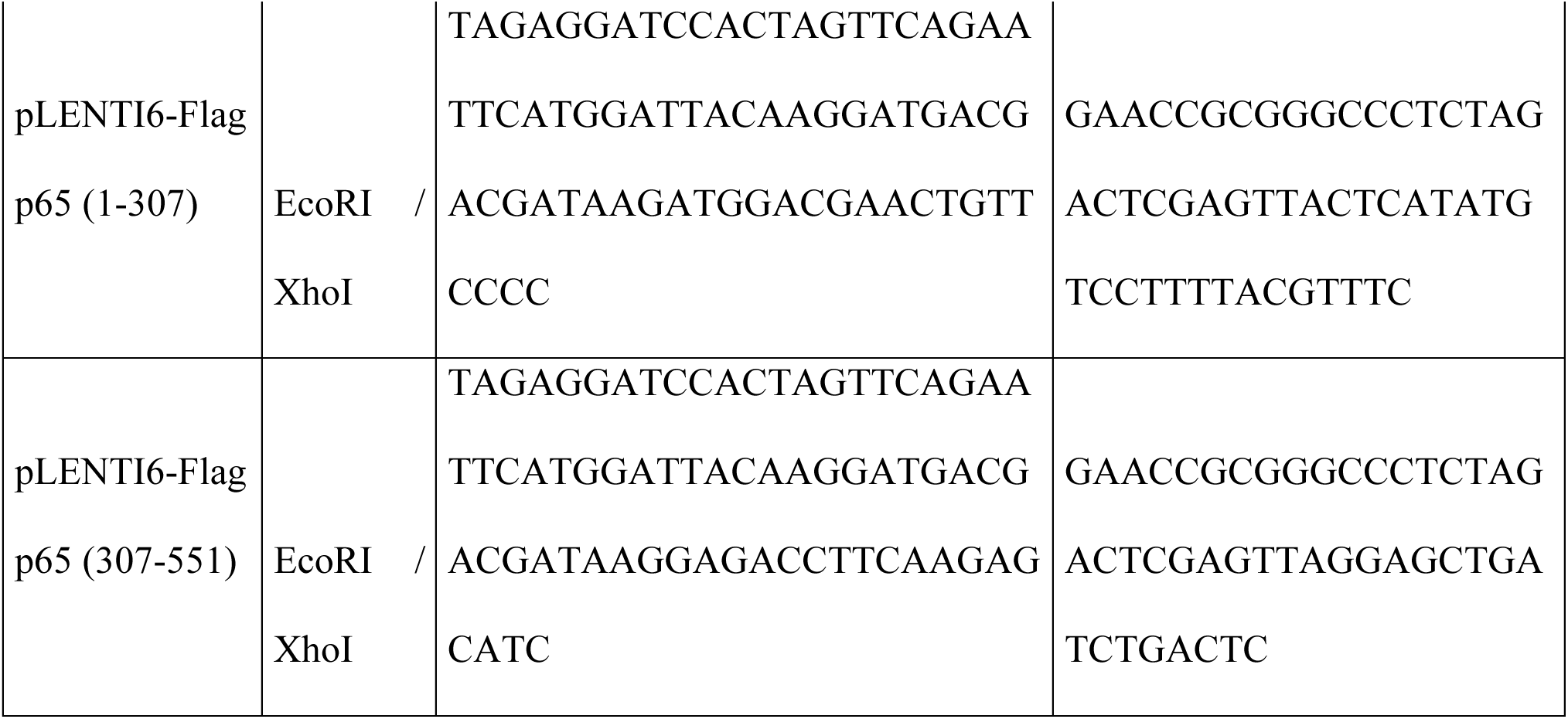

### QPCR- primer sets

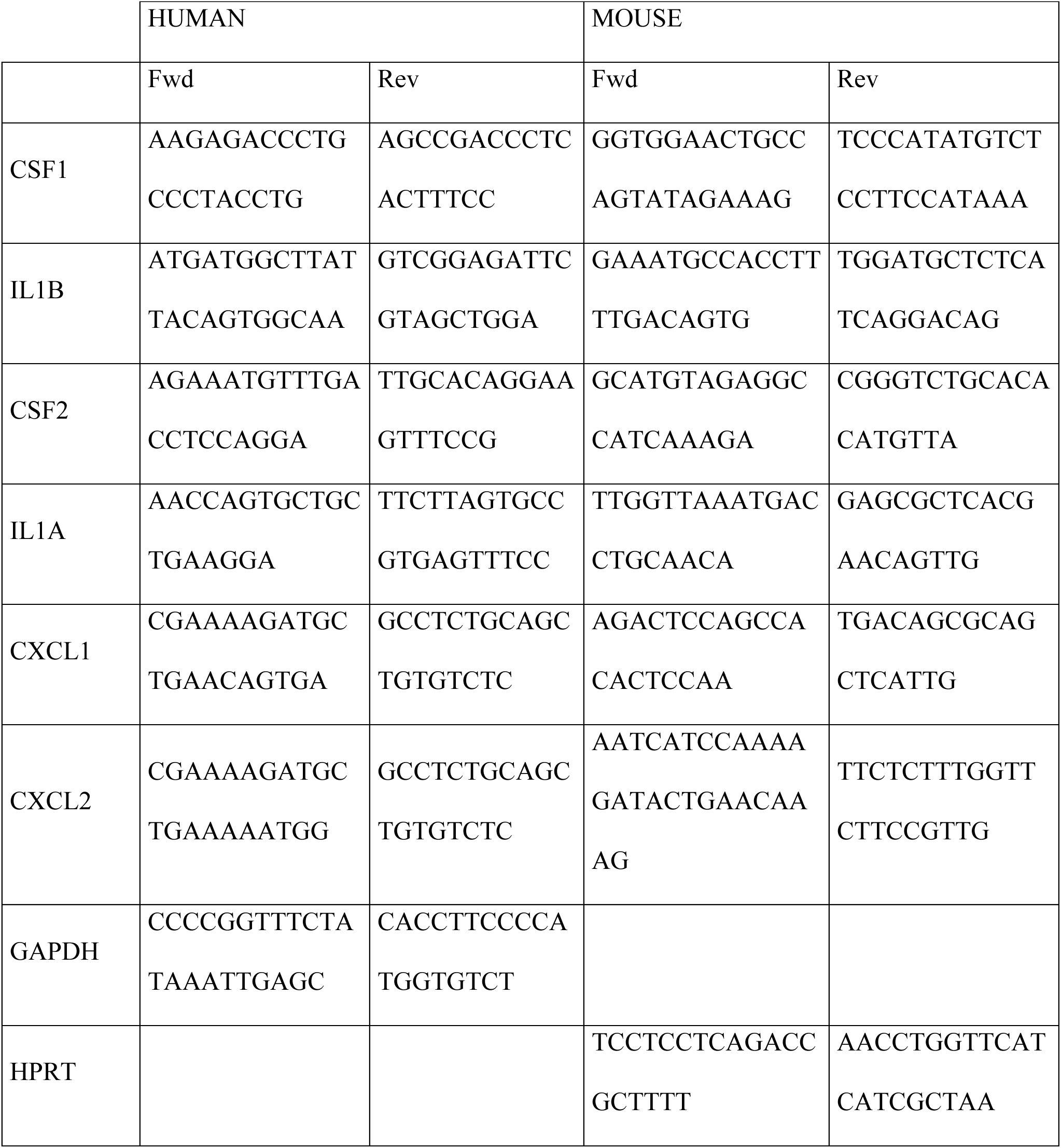

## Acknowledgements

We thank Dr Le Gallo Matthieu (Inserm U1242, CLCC Eugène Marquis, Rennes) for his technical assistance in the measure of caspase-1 activity and purification of RNAs for transcriptomic analyses. This work was supported by INCa PLBIO (PLBIO 2018-132), Ligue Contre le Cancer, Fondation ARC, Fondation de France (Price Jean Valade), and ANR PRCE (ANR-17-CE15-0027), and NIH grant R35CA197450 to M.E.P.

## Author contributions

J-P.G. and P.L. planned the experiments, performed experiments or analyzed data.

C.G., E.C-J. performed experiments or analyzed data.

M.P. and P.L. analyzed data and wrote the manuscript.

## Expanded View Figure legends

**Figure EV1.**
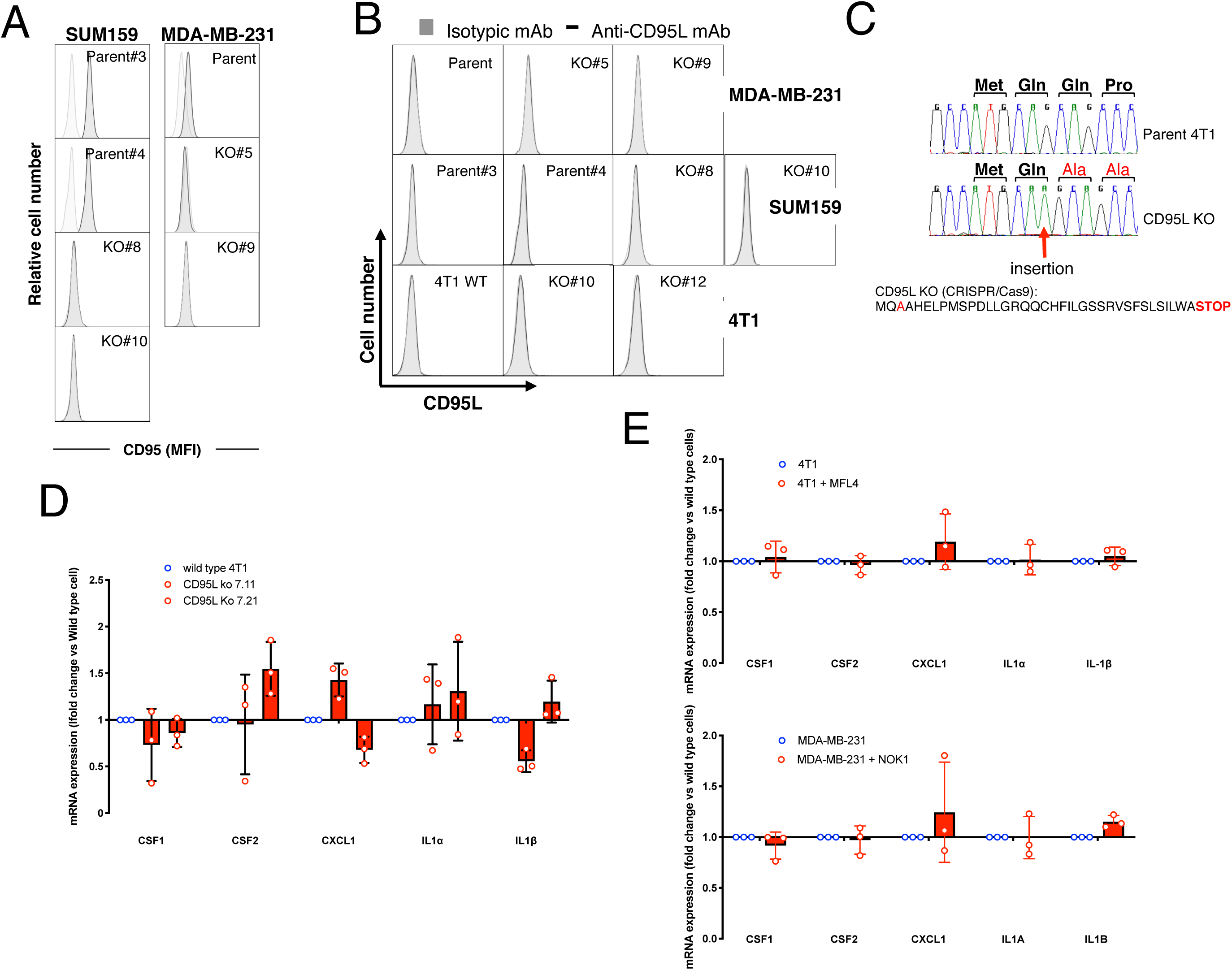
Generation and characterization of CD95-KO TNBC cells. **A.** Using flow cytometry, CD95 expression was assessed in wild type parental TNBC cells (SUM-159 and MDA-MB-231 cells) and their counterparts in which CD95 was knocked-out using CRISPR/Cas9 method. Light grey histograms represent the isotypic control staining. **B.** CD95L expression was assessed by flow cytometry in indicated cells. **C.** CRISPR/Cas9 was used to generate CD95L clones in parental 4T1 cells. Cells were cloned and CD95L DNA was sequenced. An alanine was inserted in all our CD95L-KO 4T1 TNBC cells creating a frameshift mutation that leads to a shortened polypeptide. **D.** inflammatory-gene signature expression was quantified in WT and KO-CD95L 4T1 cells using qPCR. Data are represented as gene expression (fold change) compared to wild type cells and represent mean and SD of three independently performed experiments. **E.** Inflammatory-gene signature expression was quantified in 4T1 (mouse) or MDA-MB-231 (human) WT cells incubated in the presence or absence of the neutralizing anti-mouse CD95L mAb MFL4 (*upper panel*) and the neutralizing anti-human CD95L mAb NOK1 (*upper panel*), respectively using qPCR. Data are represented as gene expression (fold change) compared to wild type cells and represent mean and SD of three independently performed experiments.

**Figure EV2.**
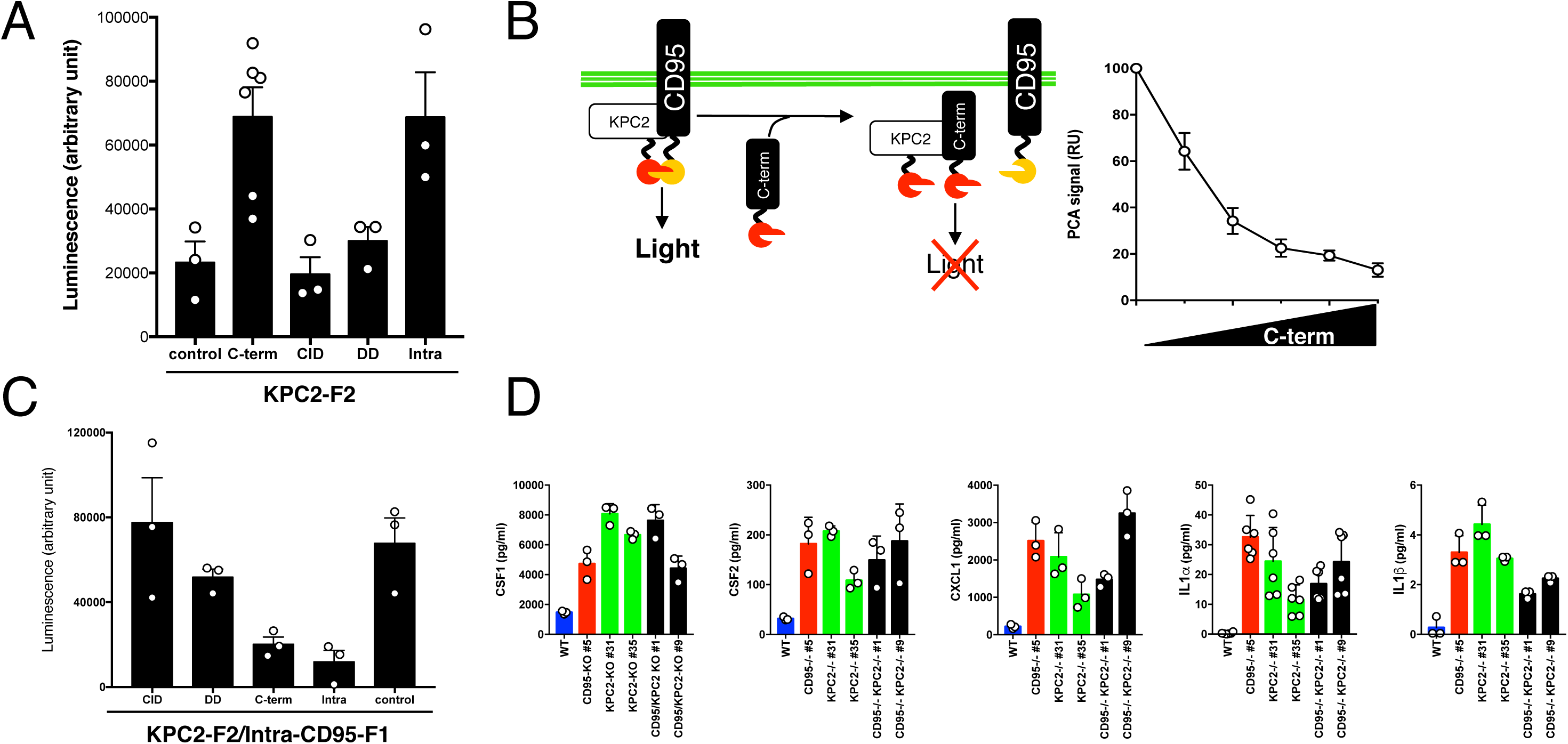
KPC2 directly interacts with the C-terminal region of CD95. **A.** HEK293T cells were co-transfected with plasmids encoding KPC2 and CD95 domains fused to the F2 domain and with the F1 fragment of Renilla Luciferase, respectively and after 24 hours, luminescence was assessed. **B.** *Left panel:* Schematic representation of the PCA competitive assay. *Right panel:* HEK293T cells were co-transfected with three plasmids encoding KPC2 and CD95 domains fused to the F2 domain and with the F1 fragment of Renilla Luciferase, respectively and C-term CD95-F2 transfected in dose-effect and after 24 hours, luminescence was assessed. **C.** HEK293T cells were transfected with plasmids encoding KPC2-F2 and the whole intracellular region of CD95 (175-319)-F1. A PCA competitive assay was developed by co-transfecting indicated F2-fused proteins and the inhibition of luminescence was assessed. Data represent mean and SD of three independently performed experiments. **D.** Inflammatory cytokines were quantified by ELISA in indicated cells and compared to parental MDA-MB-231 cells. Data represent mean and SD of three independently performed experiments.

